# Understanding biological ageing in terms of constitutive signals: *Convergence to an average decrease in cellular sensitivity and information transmission*

**DOI:** 10.1101/753566

**Authors:** Alvaro Martinez Guimera, Daryl P. Shanley

## Abstract

Biological ageing is a process that encompasses observations often too heterogeneous to draw coherent conceptual frameworks that may shed light into the generality of the underlying gradual loss of function. Whilst the concept of stochastic damage is often invoked as the driver of the ageing process, this can be too abstract to understand ageing at a higher mechanistic resolution. However, there do exist general mechanisms that describe how stochastic damage interferes with biological function, such as through genetic mutations. In a similar manner, we argue that a ‘molecular habituation’ phenomenon occurs during biological ageing where constitutive signals arising from damage accumulation drive an average decrease in network sensitivity and information transmission, as well as an increase in noise, across cells and tissue.

## 1. Introduction

A critical question at the core of research in both ageing and disease concerns how biological systems transition from a state of robust and controlled homeostasis to a state of dysregulation. In the case of biological ageing, theory points towards the abstract concept of stochastic damage as the underlying cause [1,2]. Such abstract causality seems necessary to capture the heterogeneous nature of the ageing process but is simply too vague to understand ageing at a higher mechanistic resolution.

It would be of great value to identify consistent means by which damage affects a biological system and thus provide a higher resolution into the biochemical context in which ageing takes place. One of such phenomena appears recurrently in biogerontology literature reporting dysfunctional signalling in the form of ‘blunted’ or ‘dampened’ signalling responses in cells. This can be exemplified by mTOR signalling [3,4], Nrf2 [5] and redox signalling in general [6–8], p38 signalling [9,10], HSF1 activation [11,12] and others [13–17].

We suggest that this represents a general way in which the accumulation of stochastic damage with age converges to the same effect in different signalling pathways in different biological settings.

This work discusses the possibility of this being the case and presents an analysis on the effects of constitutive signals in randomly generated biological networks that alludes to such a hypothesis.

## 2. Methods

An algorithm was developed in Matlab (MathWorks Inc., Natick, MA, 2018) for the purpose of simulating and analysing randomly generated biological networks consisting of reversible enzymatic reactions. For each algorithm settings, 10,000 different random networks were generated and a varying number of them were taken forward for further analysis. This varying number depends on whether the generated networks were responsive to a stimulus in a similar manner to biological systems (i.e. with well-defined peaks/troughs).

The generated networks can have a scale-free or random topology. Network species exist in two states: active and inactive, hence the number of nodes in the networks is double the number of defined molecular species. All network interactions follow mass-action kinetics with randomly assigned but constrained parameter values. The algorithm workflow is described in the supplementary material. The Matlab scripts are available online (https://github.com/amguimera/Random-generation-of-biological-networks).

## 3. Results

### 3.1 Rationale

Signalling pathways exist within cells as part of a large molecular interaction network. Such a network can be argued to have the following three properties:

i. Its dysregulation can give rise to constitutive signals.
ii. It is highly interconnected.
iii. It is unlikely to display ‘perfect adaptation’.

The rationale following property i) is as follows. Since homeostasis is defined as a property where an internal state can be maintained in spite of perturbations then if homeostasis fails there should be a constitutive elevation or decrease in the level of biological entities involved in the system. These are changes recognisable by baseline measurements. Constitutively elevated or diminished biological entities will be likely to serve as inputs or signals to other cellular pathways through cross-talk. In such a way, stochastic damage imprints as a substantial homeostatic perturbation to the system in the form of a constitutive signal.

Property ii) suggests that these constitutive signals are unlikely to be contained within biological pathways. Biological networks are characterised by a high degree of cross-talk between signalling pathways which allows the integration of complex signals and result in rich information processing capabilities [18–20]. However, such crosstalk between pathways can also be a source of vulnerability since they can spread the dysregulation from one pathway to another [14,21].

Property iii) gives an indication on the ability of biological networks to adapt to such constitutive signals when they spread throughout the molecular network through crosstalk. In order for a signalling system to retain a homeostatic state under the presence of a constant signal in the environment it needs to display ‘perfect adaptation’ [22,23]. This is a phenomenon where the signalling system is able to return its sensing to its original baseline levels due to the strong influence of a negative regulatory molecule that counteracts the constant stimulus from the signal.

The problem is that even within simple molecular networks of three molecular elements, where the stimulus affects the most upstream element in the signalling circuit, perfect adaptation to constitutive signals only occurs for very constrained network topologies and parameter values [24]. In the case of a large molecular interaction network, it is difficult to envision how ‘perfect adaptation’ could take place. This is because constitutive signals can affect any given pathway at a point downstream of where the negative regulator might have evolved to act upon. ‘Perfect adaptation’ at the scale of a whole-cell molecular interaction network seems unlikely to occur.

It is difficult to envision at the cellular network level how ‘perfect adaptation’ might function or evolve. One possibility is that biological pathways containing elements displaying ultrasensitive behaviour [25] might be able to buffer the effect of constitutive signals if the latter are below the switching threshold. However in this context the ultrasensitive behaviour could be activated stochastically with a greater frequency since constitutive signals might place basal levels closer to the response threshold. Without perfect adaptation, a constitutive signal in the network results in a change in the homeostatic state of the affected regulatory pathways.

Given that constitutive signals undoubtedly arise, what is the most likely consequence of a constitutive signal being present within a regulatory circuit? Should the signal be inhibitory, a given affected pathway would be rendered less responsive. Should the signal be activatory, the pathway could still lose sensitivity through the submaximal upregulation of negative regulator molecules that are insufficient to provide the system with a ‘perfect adaptation’ but render the pathway less sensitive to subsequent stimuli [26].

As an example, consider the TGFβ signalling pathway. Should there be an age-related increase in phospho-Erk levels, this would be expected to feed into the TGFβ pathway as a constitutive inhibitory signal that dampens the activation of the pathway [27]. Conversely, if phospho-Erk levels decrease with age then the TGFβ pathway could enter a state of low-level basal activation which results in in increased degradation of TGFβ receptors by I-Smad/Smurf as a negative feedback loop [28,29] and consequently a reduced responsiveness of the pathway to TGFβ stimulation.

### 3.2 Systematic tendencies in simulated biological networks

The presented rationale would suggest that biological networks have a certain tendency to lose sensitivity when constitutive signals arise. The systematic generation of *in silico* ordinary differential equation (ODE)-based models of biological networks [30,31] hint at this potentially being the case *in vivo*. A systematic approach works as follows. When any of the randomly generated models displays a realistic response to an input stimulus (in the form of a minimum number of peaks and troughs of a minimum size), they are selected for further analysis. The analysed models undergo a local sensitivity analysis for all of their kinetic parameters to systematically test how perturbations at every point in the network affects the original response to the input signal [32]. The perturbation of rate constants thus models the introduction of a constitutive signal at that point in the network, since ultimately the effect of a constitutive signal would be a change in the rate of a given reaction it is involved in. Changes in the absolute value or fold change of peaks and troughs involved in the network response to an acute stimulus are recorded and averaged for each parameter perturbation in the sensitivity analysis. Figure 1 shows representative output of a systematic workflow in terms of the structure, simulation and analysis of a randomly generated biological network with a scale-free topology [31].

**Figure 1.**
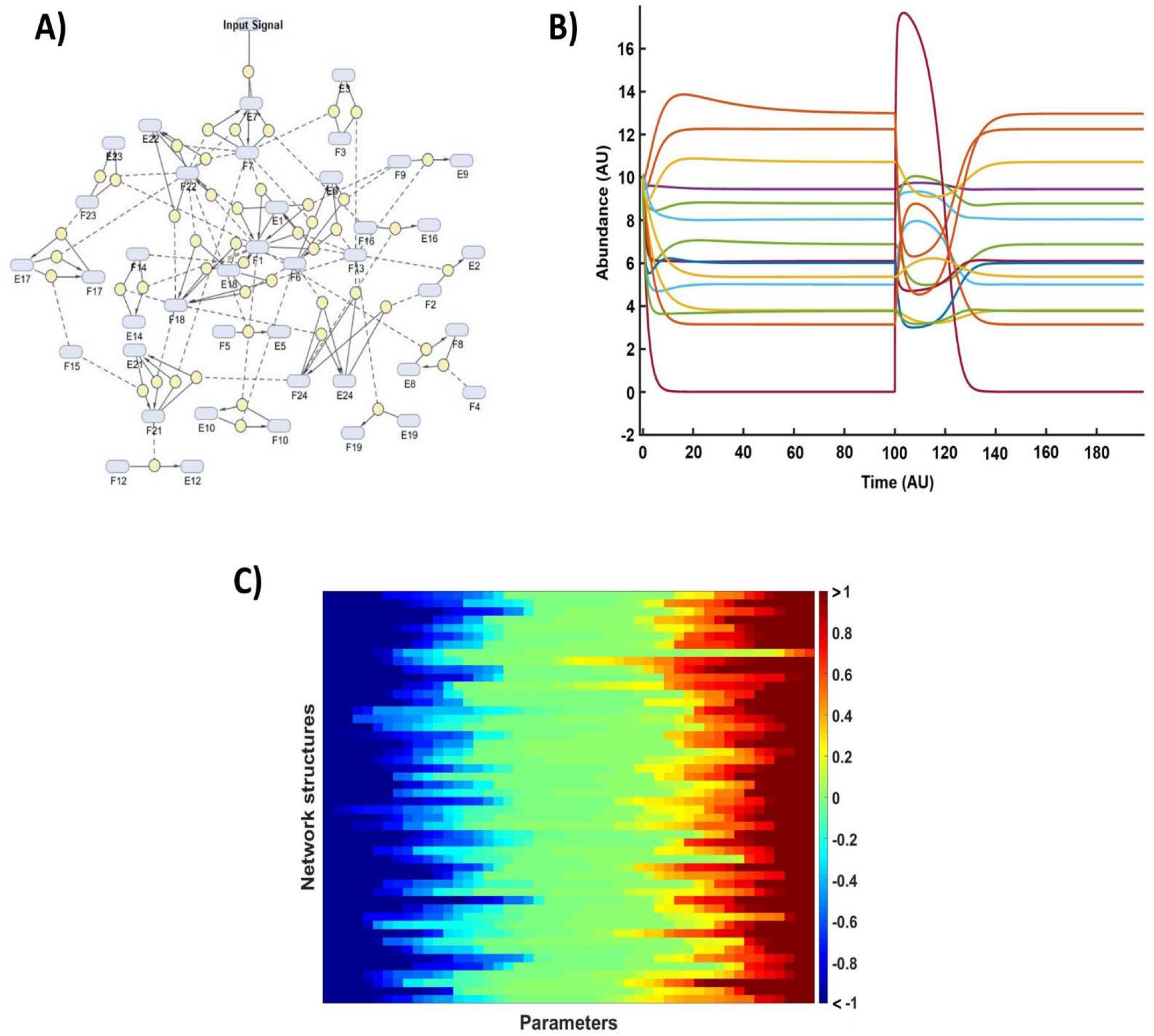
Representative algorithm output for different stages in the workflow. **A)** Generation. A biological network structure is randomly generated with certain topological features (i.e. scale-free or random). **B)** Simulation. The response of the randomly generated network to an input signal of a given strength is simulated. **C)** Analysis. If enough network elements are found to be responsive to the input signal then sensitivity analysis is performed. Colour-bar represents the average absolute- or fold-change in the response profiles such as those shown in (B).

When 10,000 scale-free biological networks consisting of 25 species and 48 interactions are generated, only ~0.5% satisfy the stimulus-responsiveness requirement (≥10 network species with a responsiveness ≥1a.u) to be taken forwards for further analysis. Sensitivity analysis on the selected models reveals there is a slight tendency for a constitutive signal to result in an average decrease in the responsiveness of network elements when assessed through the absolute levels reached upon stimulation (Figure 2A). A substantial number of parameters in the network are expected to be peripheral and not very influential in the overall dynamics of the stimulated nodes. This means they might dilute out a stronger tendency for more influential parameters. When model responses are assessed for parameters influencing the network response above a certain threshold the tendency to a loss in sensitivity is slightly more apparent (Figure 2B). These slight tendencies are also seen in networks with a different average number of interactions between nodes in the network and are also seen in random network structures that do not display a scale-free topology [31] (Supplementary Figures 1–3).

**Figure 2.**
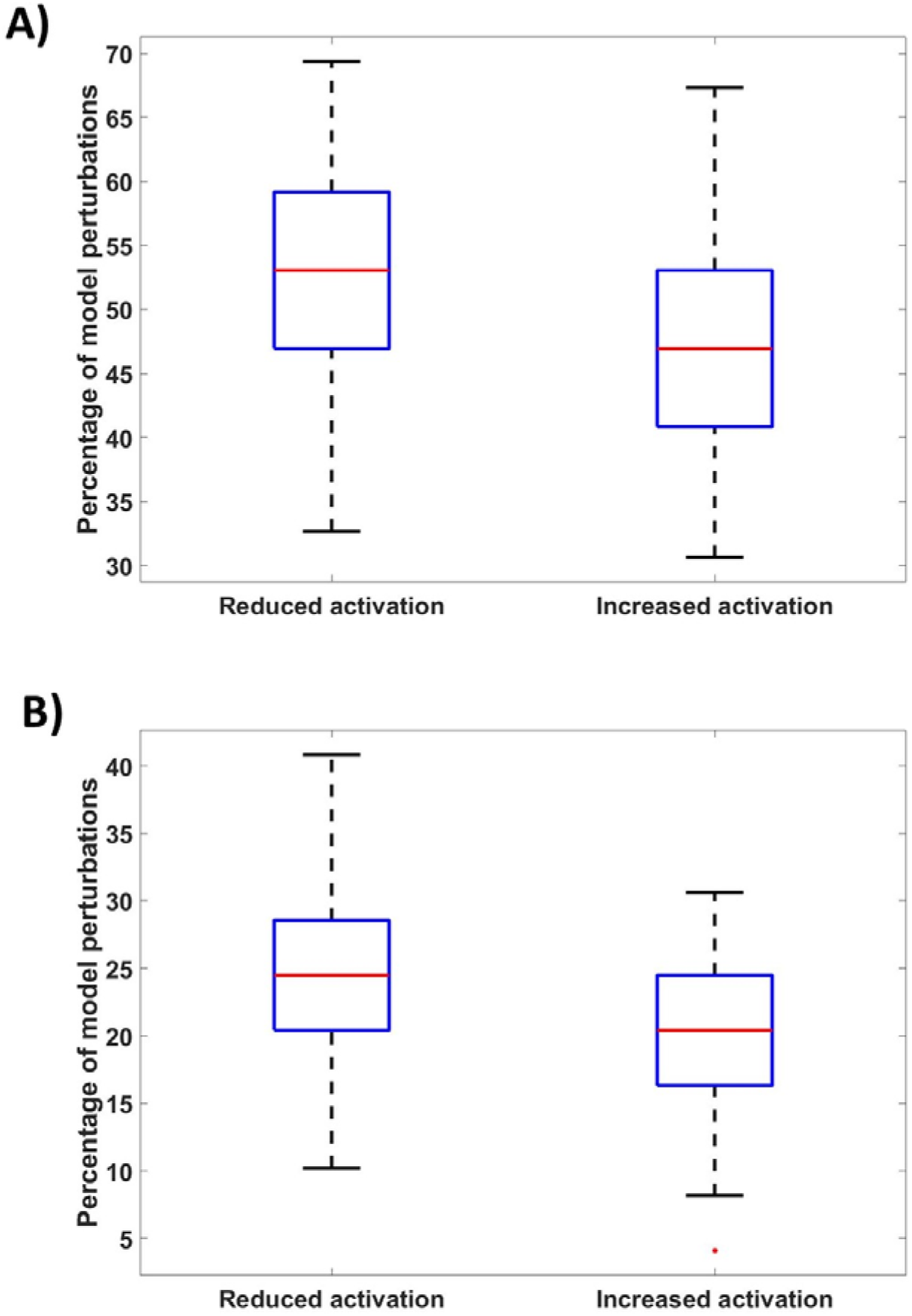
Distributions of the percentage of network parameter perturbations that result in an average decrease or increase in network responsiveness as defined by absolute-change to an input signal. **A)** Distributions for all perturbed parameters in all models simulated. P-value = 0.021. **B)** Distributions for parameters that resulted in an average change in network responsiveness of ≥0.5au. P-value = 0.00097. An outlier is defined as a value >1.5 times the interquartile range away from the upper or lower quartiles. Network topology is scale-free. Network size = 25 species (50 nodes) and 48 reactions. Number of networks analysed = 50.

When the analysis is repeated with the sole exception that network response is assessed as a fold-change from basal level, the tendency is even more pronounced (Figure 3). This makes sense since network elements that approach response saturation under the presence of constitutive signals could have no change or even an increase in the absolute levels of the responding entities alongside a decrease in their fold change from basal levels.

**Figure 3.**
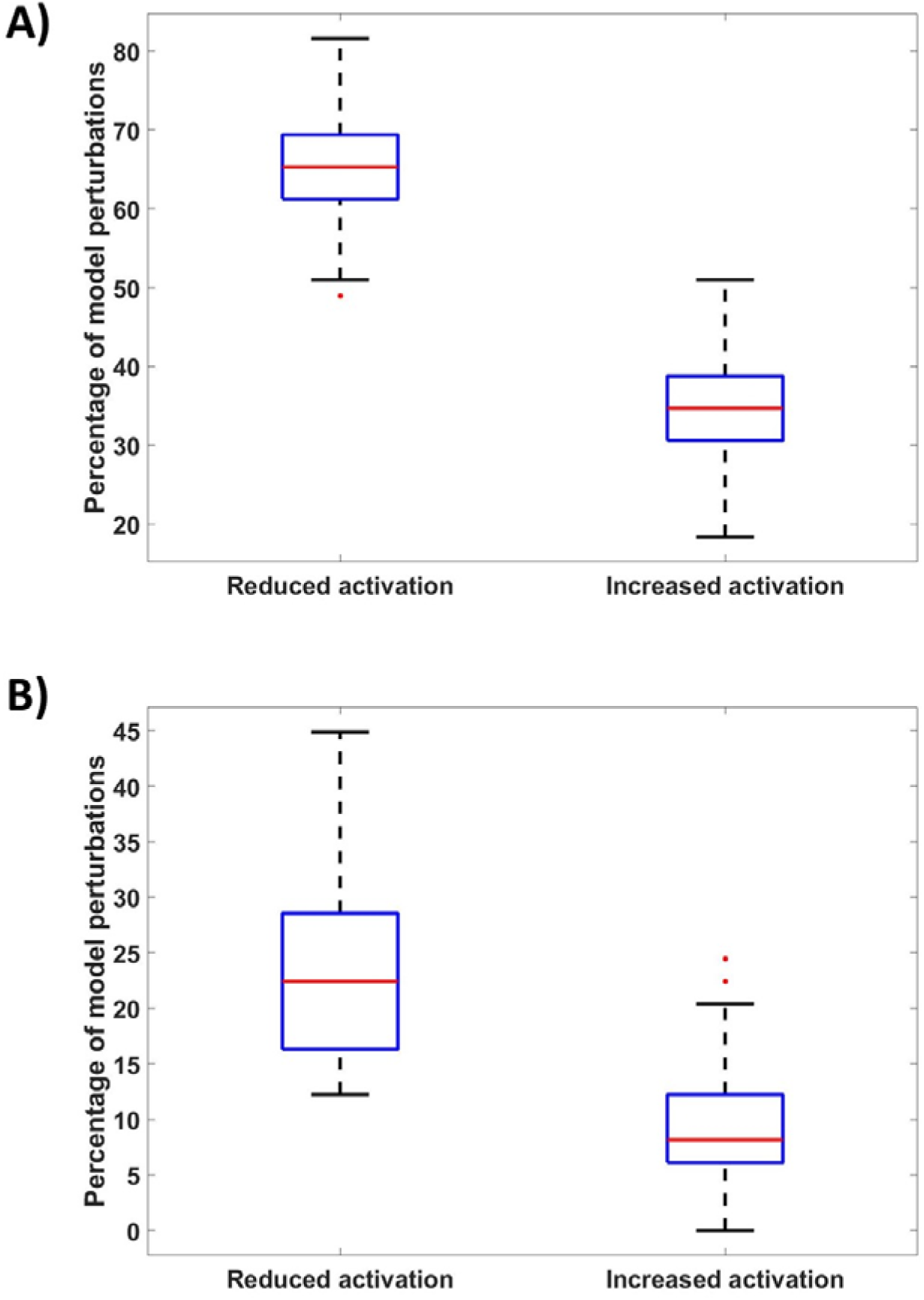
Distributions of the percentage of network parameter perturbations that result in an average decrease or increase in network responsiveness as defined by fold-change to an input signal. **A)** Distributions for all perturbed parameters in all models simulated. P-value = 7.39e^−37^. **B)** Distributions for parameters that resulted in an average change in network responsiveness of ≥0.5au. P-value = 7.39e^−17^. An outlier is defined as a value >1.5 times the interquartile range away from the upper or lower quartiles. Network topology is scale-free. Network size = 25 species (50 nodes) and 48 reactions. Number of networks analysed = 48.

Although randomly generated networks had only a slight bias towards a loss of sensitivity when assessing absolute molecule numbers, such networks are barely optimised as signal-integrating networks. Biological networks have been shaped by evolution to provide optimal responses through fine-tuned interactions. Hence, it would be expected that in the *in vivo* highly-optimised systems the bias towards a loss of sensitivity might be higher.

Such bias of constitutive signal perturbations towards causing a lower network sensitivity is also seen in a published model of ERK signalling that had its structure and kinetic parameters optimised through a parameter estimation procedure using experimental data [33]. Interestingly, a tendency towards a lower sensitivity is observed when all of the parameters are examined together but not when the most influential parameters are analysed. An assessment of all parameter perturbations regardless of magnitude of effect results in 64% of perturbations resulting in a reduced network response as judged by absolute levels reached by network entities. This is compared to 28% of perturbations that increased network response to a stimulus. These proportions become 2.38% and 9.52% respectively when only parameters having an average effect of more than 0.5 units in the network response are considered. Similar proportions are seen when responsiveness is assessed through fold-change. This emphasises the context-dependency of the resulting effect of constitutive signals on network properties such as element abundances, kinetic rates and connectivity.

## 4. Discussion

Constitutive signals can, in-principle and *in-silico*, give rise to loss of signalling effectiveness and are likely a general feature of biological ageing. If a regulatory pathway is less sensitive to a stimulus, then the distribution of abundance values that any given molecule in the pathway may have under the presence of a signal will have a greater overlap with the distribution of values that the same molecule may have under the presence of no signal (see Figure 4). Therefore, a cell is less able to discern whether the signal is in the environment or not. This translates into a greater fraction of cells within a tissue responding inappropriately to a given stimulus [26,34] and will be expected to prime for damage accumulation and a gradual loss of function.

**Figure 4.**
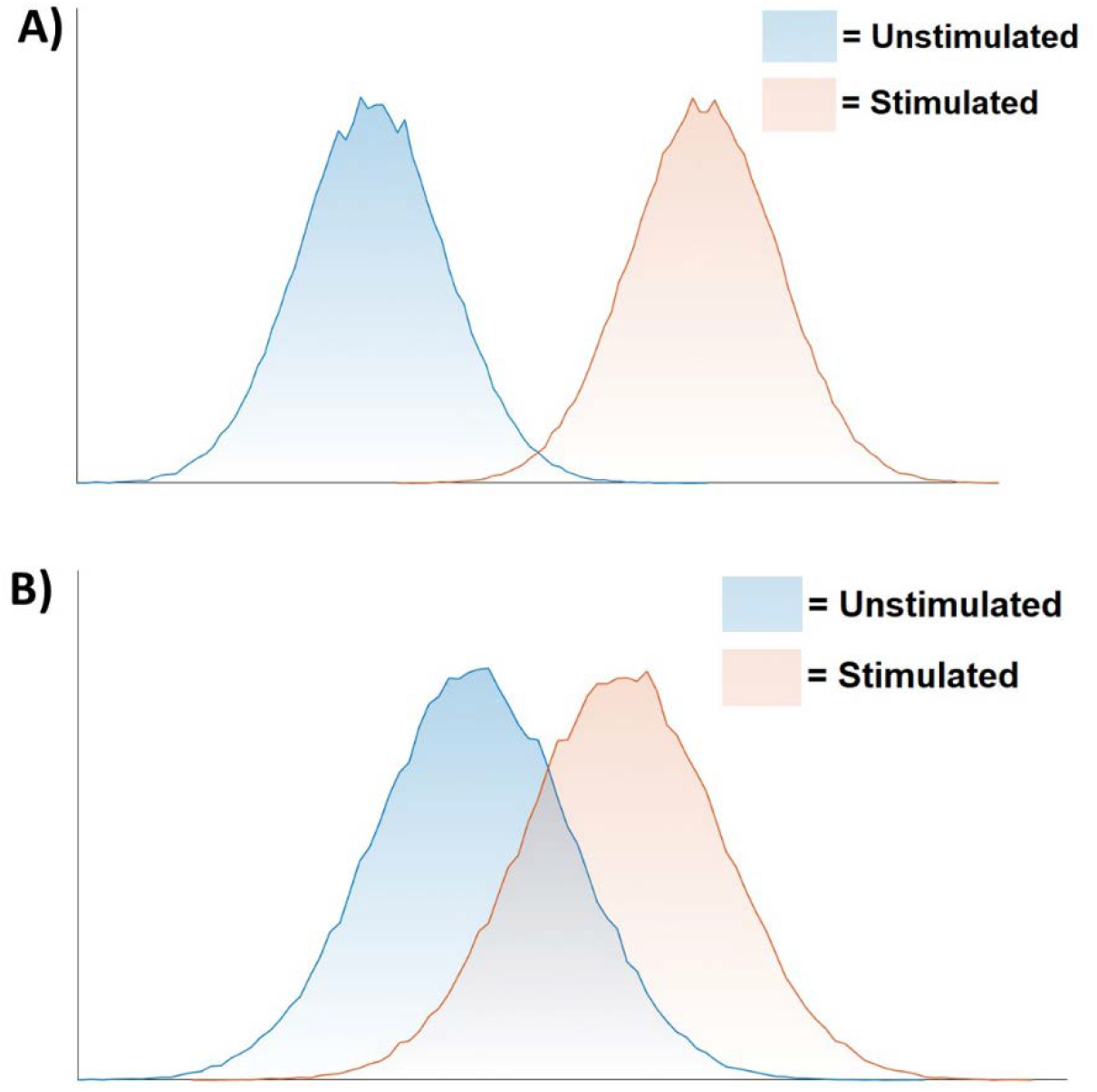
Cellular discernment of a physiological signal under **A)** healthy conditions **B)** aged conditions. A reduction in the change of the mean species abundance caused by a stimulus (loss of sensitivity) and an increase in the variance of possible abundances (increase in noise) will result in a greater overlap between the (un)stimulated distributions. Hence, cells are less able to discern the presence of the signal and this translates into a higher proportion of cells within a tissue that respond inappropriately to their environment.

The presence of constitutive signals in the form of basally elevated network activities have been shown to be a source of noise in a cellular system and to result in an overall loss of sensitivity and information transmission [26,35,36]. This is perhaps unsurprising when considering that signalling pathways have evolved to provide a close-to-optimal level of information transmission [34,37] and thus any change to the underlying parameters of the network is more likely than not to move the system away from the response optimality and reduce information transmission. Ageing would thus result in a dissipation of the rich patterns of information flow across biological networks [19,20].

The proposed sequence of events could thus involve the translation of stochastic cellular damage into permanent imprints into the system (constitutive signals) through mechanisms such a mutation or stochastically-induced self-sustaining processes [38–44]. The effect of constitutive signals would then spread throughout the molecular interaction network through crosstalk between regulatory circuits, resulting in an average decrease in pathway sensitivity and information flow and an average increase in system noise. Of course, this phenomenon would be expected to occur alongside other (perhaps more irreversible) processes that also result in a loss of network sensitivity such as altered binding affinities due to genetic mutations (Figure 5).

**Figure 5.**
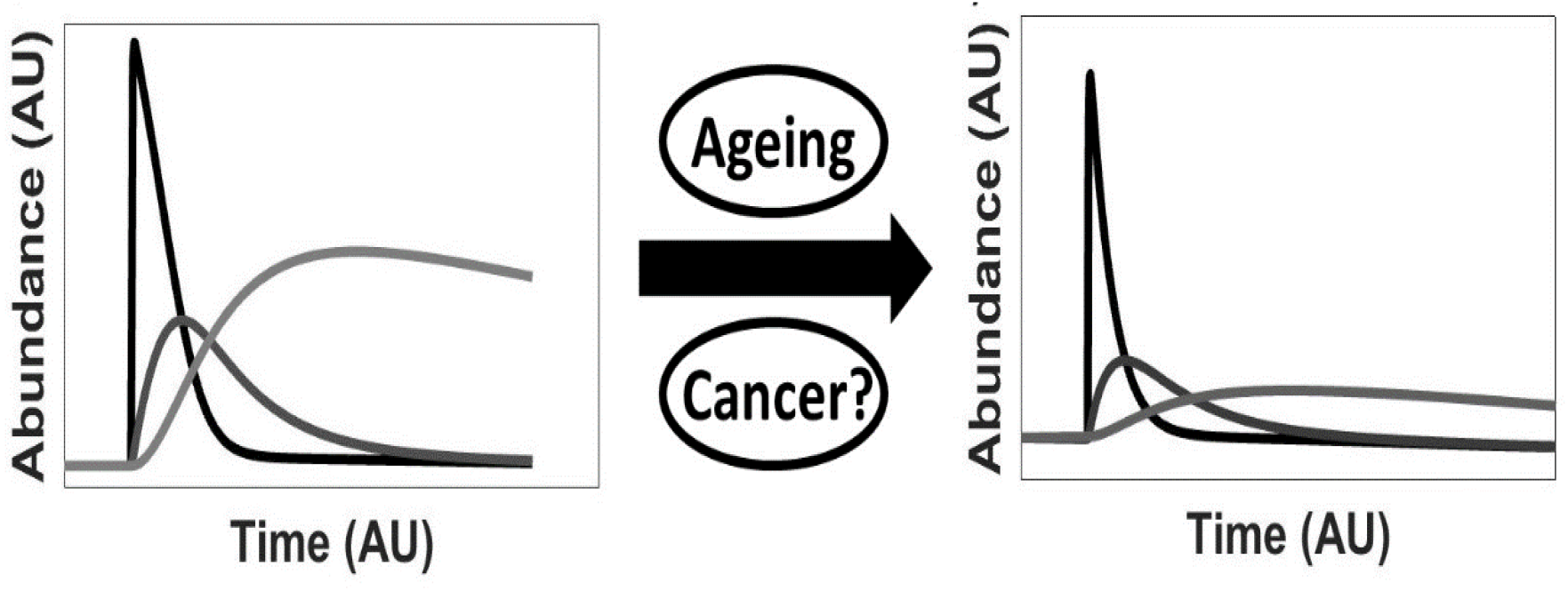
Phenomenology of changes in cellular responses with ageing and potentially cancer.

Heterogeneity in this setting could arise at various points such as the stochasticity of the transient insults (damage) that trigger the constitutive signals, the nature and the number of the constitutive signals and the wiring and parameter values characterising different affected regulatory pathways in different types of cells. In the latter case, different biological systems might contain different basal abundances of network entities or interactions of different strengths and speeds that might naturally buffer the constitutive signal and limit its percolation through certain network branches. Indeed, some pathways might be affected by constitutive signals and some might not. Hence, the phenomenon can only be described to occur on average across the cell’s entire molecular network.

In this context lifespan-extending interventions, such as NAD supplementation, might provide a local rescue in the network function. For NAD supplementation, it would be by alleviating the effects of a higher NAD degradation caused by the higher CD38 and/or PARP1 activities that are actively maintained by constitutive signals arising from chronic inflammation and/or genomic instability respectively [45,46]. However, there would still be a significant amount of dysfunctionality being actively maintained by other or the same constitutive signals in different parts of the cellular network. Constitutive signals could be stabilising a given ageing phenotype.

In physiology, the diminishing response under conditions of frequent stimulation is termed ‘habituation’. The phenomenon described here can thus be regarded as a whole cell ‘molecular habituation’ effect where ageing is characterised by an average decrease in both cell sensitivity and information transmission and an average increase in network noise. It is worth noting that information in this context represents the ability of a given entity in the cell to inform –or reduce the uncertainty-about the state of another entity in the network (the mutual information between any two species in the molecular network [26]). Such hallmarks have already been observed in senescent cells and associated with reduced intervention effectiveness [47].

It would be expected that because each cell or cell-type will have stochastic damage translate into different constitutive signals percolating through the network in different ways, that they will have distinct dysregulation ‘*omic*’ signatures. In addition, because such a ‘molecular habituation’ phenomenon presumably arises from fundamental network features and homeostatic disruption, it could arise in disease as well as biological aging. Cancer, for example, can be viewed as a disease characterised by the presence of constitutive signals [48–50].

A possible approach to test the ‘molecular habituation’ hypothesis would involve a long-duration exposure to an age-relevant signal. This could be the culture of cells with IL-1 or TNFα to model chronic inflammation [51], the knockout of the REV1 polymerase to model chronic genomic instability [52] or simply the use of a primary tissue from aged individuals which displays a basally elevated/downregulated entity that might be considered a signal. A source of information such as the STRING database [53] could be used to follow known protein-protein interactions of an identified signal into canonical pathways that might be affected. Each of the identified pathways should be sequentially stimulated with the appropriate ligand and the resulting activation magnitude compared to controls which do not display the constitutive signal. The ‘molecular habituation’ phenomenon predicts that the majority (but not necessarily all) of the stimulated pathways will have lost sensitivity in the systems exposed to the constitutive signals. It would be an interesting possibility to algorithmically explore points of introduction of reverse constitutive signals to restore system functionality.

## 5. Concluding remarks

It is a common observation in aged biological systems that some signalling pathways become dysfunctional, most often manifesting as a reduced responsiveness to stimuli. The nature of this age-related dysregulation can potentially be regarded as a ‘molecular habituation’ phenomenon in the cell’s molecular network. This involves stochastic damage resulting in homeostatic dysregulations which manifest as constitutive signals that percolate through the molecular interaction network stabilising an average state of increased noise, reduced sensitivity and reduced information transmission. It is important to consider that this might be the biochemical context in which interventions targeting the ageing process take place.

## Abbreviations

a.u: arbitrary units

## Acknowledgments

The authors are grateful to Thomas B. L. Kirkwood for discussions. This work was funded by the Novo Nordisk Fonden Challenge Programme: Harnessing the Power of Big Data to Address the Societal Challenge of Aging NNF17OC0027812.

## 7. Supplementary Material

### 7.1 Supplementary Figures

**Supplementary Figure 1.**
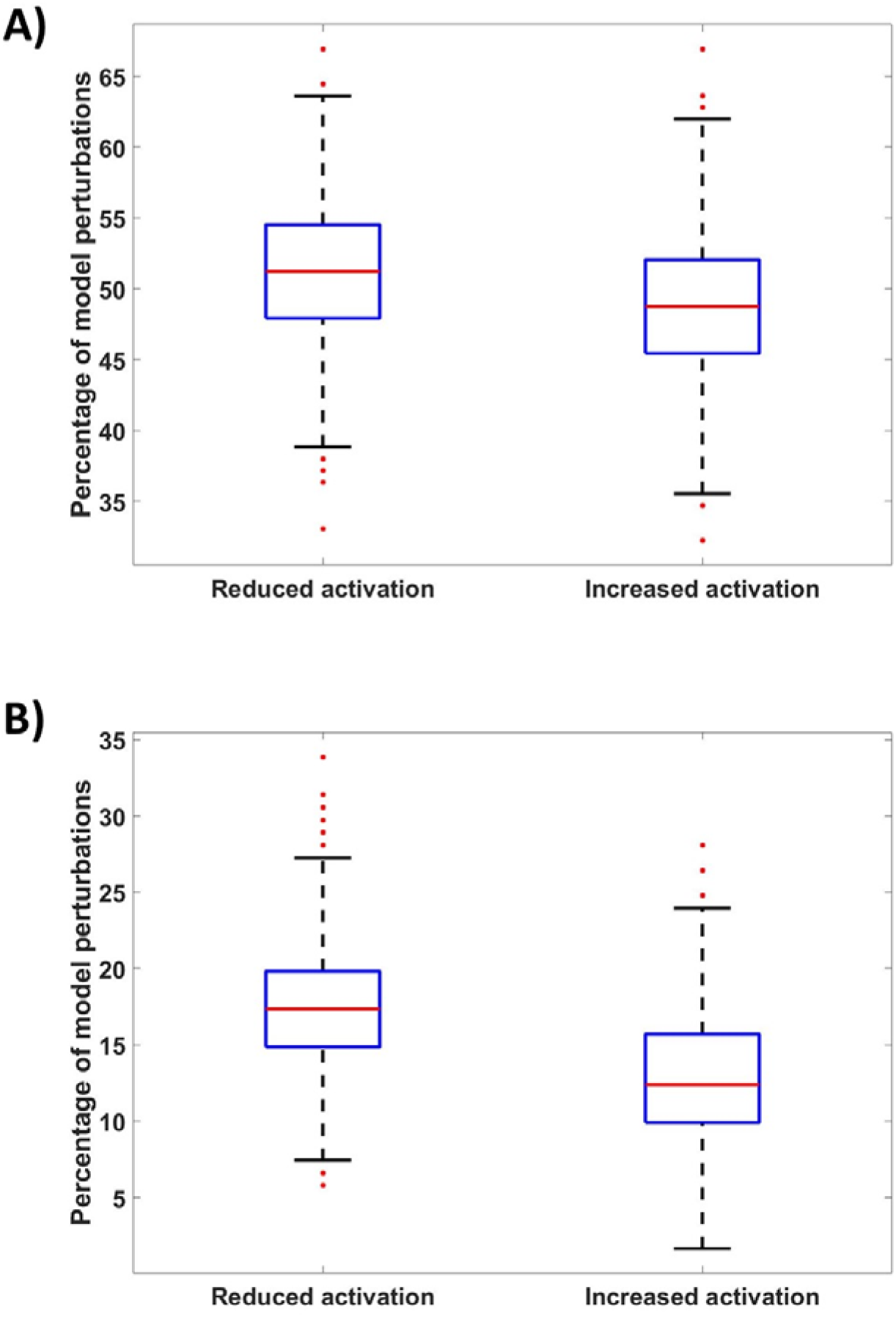
Distributions of the percentage of scale-free network parameter perturbations that result in an average decrease or increase in network responsiveness as defined by absolute-change to an input signal. **A)** Distributions for all perturbed parameters in all models simulated. P-value = 1.63e-8. **B)** Distributions for parameters that resulted in an average change in network responsiveness of ≥0.5au. 2.34e-102. An outlier is defined as a value >1.5 times the interquartile range away from the upper or lower quartiles. Network topology is scale-free. Network size = 25 species (50 nodes) and 120 reactions. Number of networks analysed = 893.

**Supplementary Figure 2.**
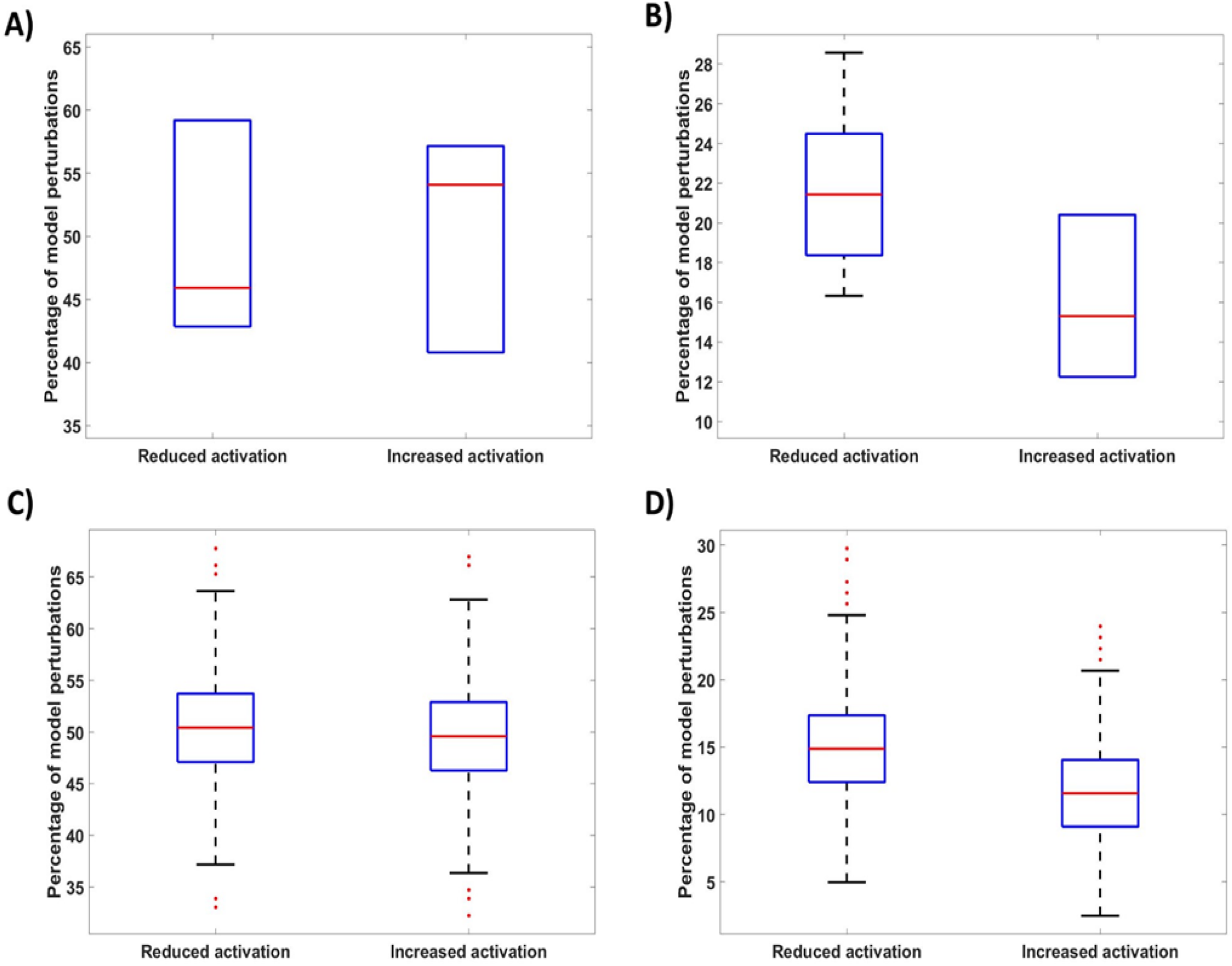
Distributions of the percentage of random-topology network parameter perturbations that result in an average decrease or increase in network responsiveness as defined by absolute-change to an input signal. **A)** Distributions for all perturbed parameters in 6 models with 25 species (50 nodes) and 48 reactions. P-value = 0.77. **B)** Distributions for parameters that resulted in an average change in network responsiveness of ≥0.5au in models with 25 species (50 nodes) and 48 reactions. P-value = 0.034.**C)** Distributions for all perturbed parameters in 820 models with 25 species (50 nodes) and 120 reactions. P-value = 1.42e-6. **D)** Distributions for parameters that resulted in an average change in network responsiveness of ≥0.5au in models with 25 species (50 nodes) and 120 reactions. P-value = 3.96e-56. An outlier is defined as a value >1.5 times the interquartile range away from the upper or lower quartiles. Network topologies are random. Number of networks analysed in A) and B) is 6. Number of networks analysed in C) and D) is 820.

**Supplementary Figure 3.**
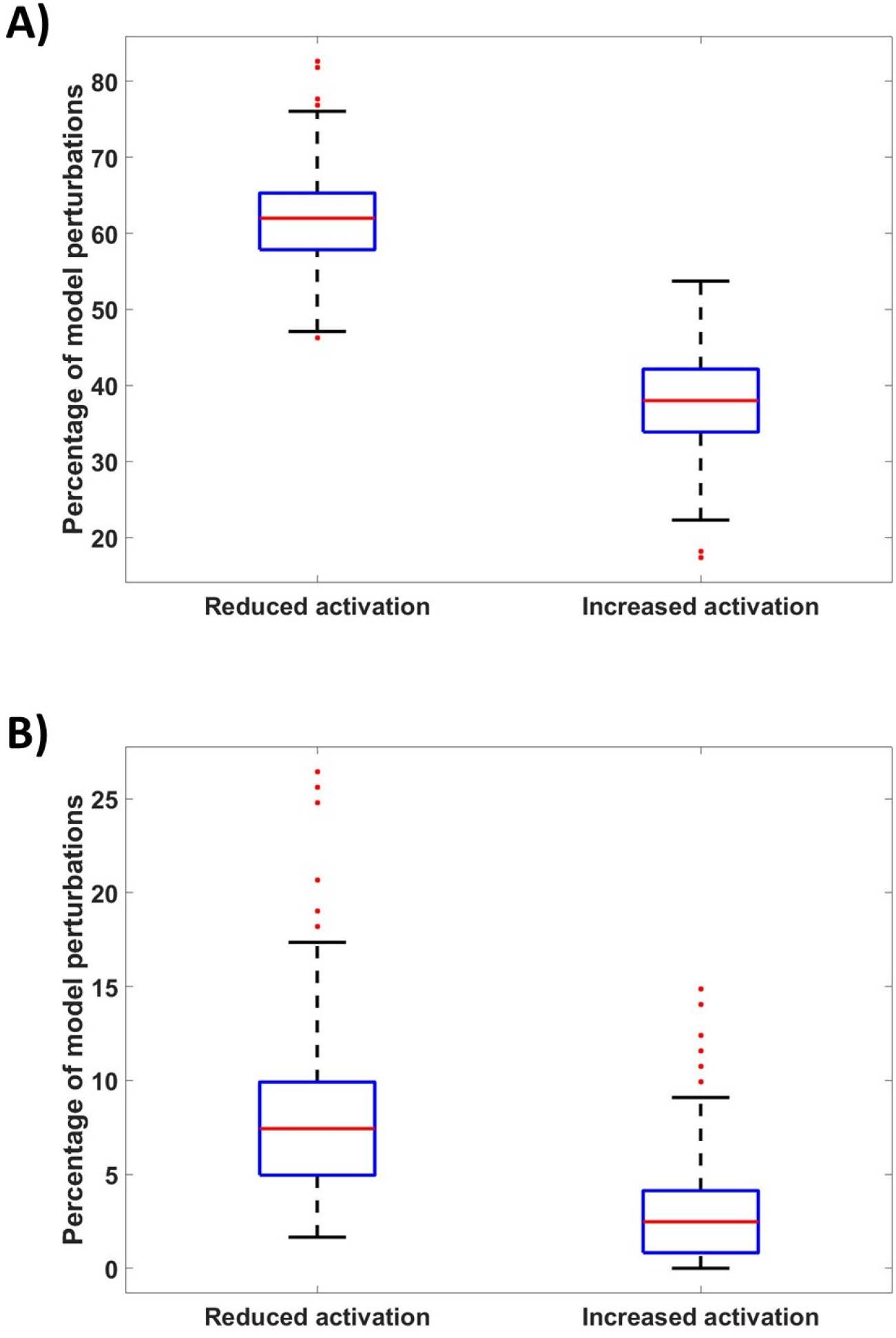
Distributions of the percentage of scale-free network parameter perturbations that result in an average decrease or increase in network responsiveness as defined by fold-change to an input signal. **A)** Distributions for all perturbed parameters in all models simulated. P-value = 0. **B)** Distributions for parameters that resulted in an average change in network responsiveness of ≥0.5au. P-value = 1.36e-188. An outlier is defined as a value >1.5 times the interquartile range away from the upper or lower quartiles. Network topology is scale-free. Network size = 25 species (50 nodes) and 120 reactions. Number of networks analysed = 852.

### 7.2 Algorithm workflow

An algorithm was developed in Matlab (MathWorks Inc., Natick, MA, 2018) for the purpose of simulating and analysing randomly generated biological networks consisting of reversible enzymatic reactions. The algorithm workflow is as follows:

i. User defines the number of species in the network as well as their uniform initial abundance (au) and scale of rate constant values.
  - Note that noise will be added to this value.
ii. The algorithm creates a numerical vector of size corresponding to the user-defined number of species so that each species has a unique numerical identifier.
iii. The species are duplicated into *E*_*n*_ species and *F*_*n*_ species where *E* corresponds to the inactive form of the species with identifier *n* and *F* corresponds to the active form of the species with identifier *n*.
iv. A random *F*_*i*_ species is chosen. The algorithm can then chose a random *E*_*n*_ species or a random *F*_*n*_ species with equal probability.
  - If an *E*_*n*_ species is chosen an activatory reaction is formalised as:

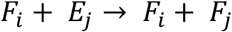 And assigned a rate constant *k* where *k*_*act*_ = *k*.
  - If an *F*_*n*_ species is chosen an inhibitory reaction is formalised as:

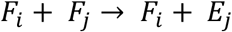 And assigned a constant *k* where *k*_*inhib*_ = 2 ⋅ *k*.
  - Note that the selection of a random *E*_*n*_or *F*_*n*_ value will follow a uniform probability in the case of random topology network or will be biased in favour of species with more existing interactions in the case of a scale-free network topology.
v. Repeat step (iv) for a user-defined number of reactions.
  - User defines the number of reactions as a proportion of the total number of all possible interactions 2*N*^2^ − 2*N* where *N* is the user-defined number of species in the network. Such interactions encompass all pairwise interactions between *F* species as well as interactions between *F* species and *E* species whilst excluding all self-interacting reactions for simplicity.
vi. The generated rate constants, initial abundances as well as the strings corresponding to reactions, rate laws and species names are inputted into ‘Simbiology’ to create an ordinary differential equation (ODE) – based model where all reactions follow mass action kinetics. An extra reaction in the form of *InputSignal* → *E*_*n*_ is introduced to affect a randomly chosen *E*_*n*_ species to model a network stimulus.
  - This stimulus is introduced as an ‘Event’ after time is allowed for model equilibration. Default stimulus strength = 10⋅InitialSpeciesAbundance.
vii. The model is simulated using Matlab’s ode15s solver. If the model displays a minimum number (default = 10) of species which display a recognisable change from steady state in response to the network stimulus (in the form of peaks/troughs) the absolute- or fold-change values for the responding species are saved into an excel file with a unique model identifier.
  - Note that only *F*_*n*_ species are analysed for peaks/troughs.
  - The Simbiology model file is also saved with the same unique identifier.
viii. Steps (iv) to (vii) are repeated for a user-defined number of randomly generated models.
ix. *A separate script must be run in the same directory which iteratively* opens each saved model and its corresponding excel file.
x. All the parameters in each imported model are sequentially perturbed (default = x10) and the absolute- or fold-change from the corresponding reference peaks/troughs saved in the excel file are calculated and averaged.
  - An increase in peak/trough responsiveness to the stimulus will show as positive numbers.
  - A decrease in peak/trough responsiveness to the stimulus will show as negative numbers.
  - Results display the average change in the absolute- or fold-change value of responding species in the network for each parameter perturbation (x-axis) for each network structure probed (y-axis). Results can be viewed as a heatmap and saved as an excel file.

